# Functional dissociations between prefrontal and parietal cortex during task switching: A combined fMRI and TMS study

**DOI:** 10.1101/2022.12.15.520659

**Authors:** José A. Periáñez, Raquel Viejo-Sobera, Genny Lubrini, Juan Álvarez-Linera, Elisa Rodríguez Toscano, María Dolores Moreno, Celso Arango, Diego Redolar-Ripoll, Elena Muñoz Marrón, Marcos Ríos-Lago

## Abstract

Preparatory control in task-switching has been suggested to rely upon a set of distributed regions within a frontoparietal network, with frontal and parietal cortical areas cooperating to implement switch-specific preparation processes. Although recent causal evidences using transcranial magnetic stimulation (TMS) have generally supported this model, alternative evidences from both functional neuroimaging and neurophysiological studies have questioned the switch-specific role of both frontal and parietal cortices. The aim of the present study was to clarify the role of prefrontal and parietal areas supporting preparatory cognitive control in task-switching. Within this purpose, an fMRI study during task-switching performance was conducted to identify the specific brain areas involved in preparatory control during performance of a task-switching paradigm. Then, TMS was applied over the specific coordinates previously identified through fMRI, that is, the anterior portion of the inferior frontal junction (aIFJ) and the intraparietal sulcus (IPS). Results revealed that TMS over the aIFJ disrupted performance in both switch and repeat trails in terms of delayed responses as compared to Sham condition. In contrast, TMS over the IPS selectively interfered performance in switch trials. These findings support a multicomponent model of executive control with the aIFJ being involved in more general switch-unspecific process such as the episodic retrieval of goals, and the IPS being related to the implementation of switch-specific preparation mechanisms for activating stimulus-response mappings. The results help conciliating preceding evidences about the role of a frontoparietal network during task-switching, and support current models about a hierarchical organization within prefrontal cortex.

**Significance Statement:** A combined fMRI and TMS study was conducted to clarify the brain areas involved in the executive control of attention during a cueing task-switching paradigm.

Functional dissociations were observed during TMS stimulation, with prefrontal and parietal areas playing different roles during task-switching preparation. While the anterior portion of the inferior prefrontal junction seemed to be involved in a general mechanism of memory retrieval for goal identification, the intraparietal sulcus seemed to be engaged in a switch-specific mechanism for the translation of abstract task goals into action rules.

The results help conciliating preceding evidences about the role of a frontoparietal network during task-switching, and support current models about a hierarchical organization within prefrontal cortex.

## Introduction

Neuroanatomical models about executive control suggest interactions within a frontoparietal network (FPN) of brain regions cooperating for an adjusted and flexible control of human behaviour (Miller and Cohen, 2001; Koechlin and Summerfield, 2007; Petersen and Posner, 2012). General consensus exists among theories assuming that the prefrontal cortex will generate top-down signals to bias processing in posterior areas. Studies using task-switching paradigms in combination with neuroimaging, neurophysiological, and lesional techniques have contributed to clarify the role of different brain regions within this FPN to cognitive control (Sohn et al., 2000; Monchi et al., 2001; Periáñez et al., 2004; Kopp et al., 2006; Niendam et al., 2012; Muhle-Karbe et al., 2014; Worringer et al., 2019). Thus, task-switching requires to change frequently among a set of simple tasks, which is supposed to activate certain control operations that will ultimately allow a flexible adaptive behaviour (Wylie and Allport, 2000; Monsell, 2003; Logan and Bundesen, 2003).

Functional neuroimaging studies have identified switch-specific activation in prefrontal areas such as medial frontal gyrus, inferior frontal gyrus, and the inferior frontal junction (IFJ), and in parietal areas such as posterior parietal cortex and precuneus. For instance, the IFJ has been suggested to exert top-down preparatory control during switch trials for the updating of abstract task representations (Derrfuss, et al., 2005; Kim et al., 2012; Worringer et al., 2019). Within the parietal lobe, the intraparietal sulcus (IPS) would provide a more action-related task representations such as stimulus-response mappings (Andersen et al., 1997; Culham and Kanwisher, 2001; Worringer et al., 2019). The analysis of the time dynamics of the activation within this FPN by means of neurophysiological techniques have generally supported the idea that activation of prefrontal regions precedes the activation of posterior ones during the implementation of a switch in task (Stuss and Picton, 1978; Periáñez et al., 2004; Brass et al., 2005; Kopp et al., 2006).

Transcranial Magnetic Stimulation (TMS) has offered causal evidences about the contribution of different prefrontal (Rushworth et al., 2002; Muhle-Karbe et al., 2014) and parietal areas (Muhle-Karbe et al., 2014) to preparatory control in task-switching by lowering brain activity. Thus, Muhle-Karbe et al. (2014) found that the application of online inhibitory TMS over the IFJ increased reaction times in switch trials in an experimental condition that involved the updating of the abstract task rules. In addition, TMS inhibition of the IPS increased the percentage of errors when switching involved remapping stimulus-response associations. The authors interpreted that prefrontal and parietal cortices would implement switch-specific preparation processes operating at two different levels of abstraction (i.e., task-goals and response-sets). However, alternative data have questioned the switch-specific preparatory role of prefrontal and parietal brain regions during task-switching (Bode and Haynes, 2009; Mansfield et al., 2012; Ruge et al., 2013). For instance, evidence about prefrontal activation during both switch and repeat trials suggests that prefrontal areas could implement more general switch-unspecific mechanisms (Braver et al., 2003; Barceló et al., 2008; Jamadar et al., 2010). Unfortunately, no other TMS study has explored further these alternative ideas, thus leading open the question about the role of the FPN in task-switching.

The aim of this work was to clarify the role of prefrontal and parietal areas supporting preparatory control during task-switching performance in a combined fMRI and TMS study. Performance of a well-known cueing task-switching paradigm using fMRI was analysed to identify brain areas associated to preparatory task-switching; then, reducing excitability of those areas using TMS allowed to analyse its disruptive behavioural effects on this task. Following preceding results (Muhle-Karbe et al., 2014), it was hypothesized that if both prefrontal and parietal cortices implement switch-specific preparation processes, then TMS over these areas should interfere with performance in switch, but not in repeat trials. Alternatively, if the activity of either prefrontal or parietal cortices during task-switching relates to a more general switch-unspecific process, then TMS over these regions would interfere performance in both switch and repeat trials.

## Materials and methods

### Participants

A study combining fMRI and TMS techniques in two separate sessions with two different groups of participants was conducted. fMRI activation data were used to determine the most suitable target areas for TMS stimulation. Twenty healthy university students (mean age 26.8±1.3 years; 12 female) participated in the fMRI study. Each participant underwent a screening interview excluding sensory-motor problems, history of neurological, or psychiatric problems, or substance abuse. Twenty healthy university students (mean age 29.3±6.8 years; 13 female) participated in the TMS study. All of them met the TMS safety criteria (Rossi et al., 2009). They had normal, or corrected to normal, visual acuity, and none were taking any medication with effects over the nervous system, or had history of neurological or psychiatric disorder, or drug/alcohol abuse. All subjects signed informed consent for their participation. The study was approved by the Ethics Committees of both the Universitat Oberta de Catalunya, and the Gregorio Marañón University Hospital. All participants gave written informed consent to participate in the study following the Declaration of Helsinki.

### Experimental Task

The experimental task-switching paradigm was inspired by the classical numerical judgment task by Allport et al. (1994). Individuals were instructed to respond to target stimuli centred on the screen (a number between 1 and 9, excluding 5) according to one of two possible task rules: odd/even or >5/<5 (see Figure 1a). Task rules were indicated on the basis of a trial-by-trial task-cueing procedure being cues presented between 1000-2000 milliseconds prior to the target number. This long cue-to-target interval was selected to favour cue-related task preparation and associated switch-related BOLD activation to occur (Ruge et al., 2013). The cues were two different symbols presented during 300 milliseconds duration that appeared centred on the screen: “x” or “+” for the odd/even and >5/<5 tasks, respectively. All the stimuli were presented in white colour over a black background in Arial font with a physical dimension of 1 × 1 inches. The target display remained on the screen for 300 milliseconds with a response time limit of 3000 milliseconds from the target onset. Participants were instructed to respond to target numbers according to whether the number was odd (left index finger) or even (right index finger) if the preceding cue was the “x” symbol, or whether the number was < 5 (left index finger) or >5 (right index finger) if the preceding cue was the “+” symbol. Responses were immediately followed by a 100 milliseconds feedback indicating right, wrong, too fast, or too slow performance. The interval between each response and the next cue adopted a random value between 1000-3000 milliseconds (see Figure 1). Both, speed and accuracy were stressed in the written instructions presented at the beginning of the task. The experimental task was controlled by Presentation software (www.neurobs.com).

**Figure 1:**
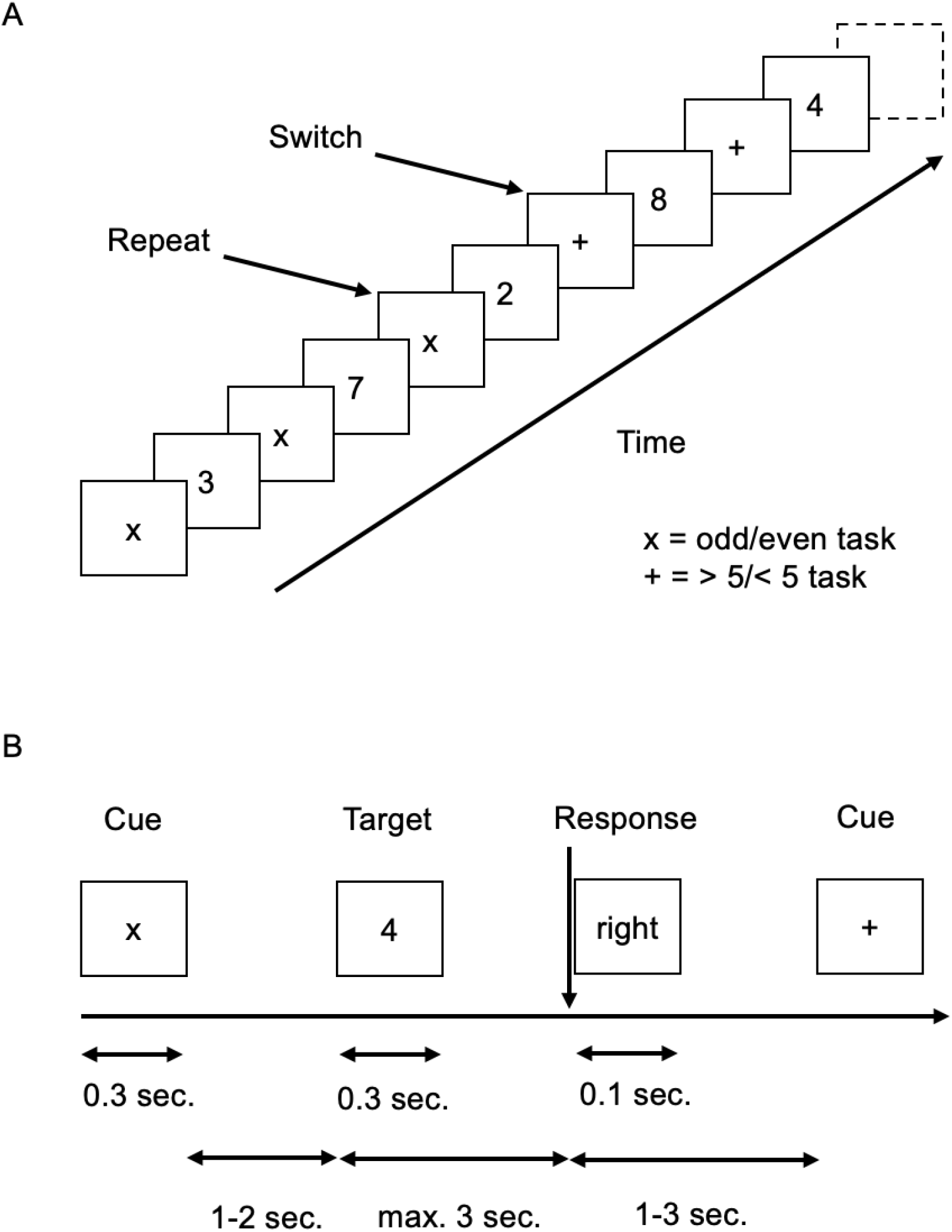
Stimulus material and experimental conditions. (a) Schematic of trial sequences illustrating the two experimental conditions being analysed. Switch trials involved a change in task rule as compared with the immediately preceding trial while repeat trials involved using the same rule as before (either odd/even task or > 5/< 5 task. (b) Each trial consisted of a visual cue (“x” or “+” symbols) followed by a visual target display with one number between 1 and 9, excluding 5. Participants were instructed to respond to target numbers according to whether the number was odd (left index finger) or even (right index finger) if the preceding cue was the “x” symbol, or whether the number was < 5 (left index finger) or >5 (right index finger) if the preceding cue was the “+” symbol. Responses were immediately followed by feedback indicating right, wrong, too fast, or too slow performance.

Participants from both fMRI and TMS experiment complete blocks of 154 trials organized semi-randomly in 42 series consisting of an initial switch trial being followed by a varying number of repeat trials (between 2 and 4). The switch condition included the initial trial of a series where the current cue indicated to perform a different task than the one performed in the preceding trial (either odd/even or >5/<5). The repeat condition included the last trial of a series (either the 2^nd^, the 3^rd^, or the 4^th^ trial after a switch trial) where the current cue indicated to respond according to the same rule as in the preceding trials. The experimental session lasted around 12 min. Before the experimental task participants practice the task until the experimenter was sure that they had understood the instructions. Reaction Times (RTs) were measured in both the first and the last correct trials of each series, corresponding with switch and repeat tasks conditions, respectively.

### Behavioural analyses

The percentage of correct responses and RTs to correct trials were analysed in the fMRI study as a function of the Task factor (switch versus repeat) by means of Student *t* tests. In addition, in the TMS study, the percentage of correct responses and RTs to correct trials were each analysed by means of a 3 × 2 repeated measure ANOVA with the factors Area of stimulation (Prefrontal, Parietal, Sham) and Task (switch versus repeat). A significance level of 0.05 was set for all contrasts. A Bonferroni-corrected significance level of p < 0.05 was adopted for all tests of simple effects involving multiple comparisons. SPSS version 22.0 statistical software package was used for all the analyses.

### fMRI scanning

The fMRI data were acquired on a 3.0T SignaHDx MR scanner (GE Healthcare, Waukesha, WI) using an eight-channel head coil (GE Coils, Cleveland, OH). Head motion was minimized with a vacuum pack system molded to fit each subject. Functional images were obtained with a gradient echoplanar sequence using blood oxygenation level–dependent (BOLD) contrast, each comprising a full volume of 39 contiguous axial slices (3 mm thickness, 0 mm spacing) covering the whole brain. Volumes were acquired continuously with a repetition time (TR) of 3 s [TE = 31 ms, flip angle = 90, field of view (FOV) = 21.7 cm]. A total of 420 scans were acquired for each participant in a single session (21 min run), with the first five volumes subsequently discarded to allow for T1 equilibration effects. High-resolution T1-weighted spoiled gradient recall (SPGR) anatomical images were also obtained for each participant (184 1.8-mm-thick axial images, TR = 5.5, TE = 2.3, FOV = 24 cm, 256x256 matrix).

The Presentation software package (https://www.neurobs.com/) was used for stimuli presentation. Stimuli were presented to participants through optic-fiber-based glasses (MRVision 2000 ultra, Resonance Technology, Inc., Northridge, USA) connected to the stimulation computer. Responses were registered with Lumina LP400 response pads for fMRI.

### fMRI data analysis

The data were analysed using a general linear model in SPM12 (Welcome Department of Imaging Neuroscience, London, UK, www.fil.ion.ucl.ac.uk/spm/) implemented in MATLAB R2018a (Mathworks, Inc., Sherborn, MA). Individual scans were i) spatially realigned and unwarped to compensate for head-movement; ii) corrected for differences in slice acquisition time (slice timing correction); iii) spatially normalized and iv) spatially smoothed to reduce noise and to compensate for anatomical inter-subject variability (Gaussian filter of 8 mm FWHM), using standard SPM methods. Population inference was made through a two-stage procedure. At the first level, we specified in a subject-specific analysis where the event-related activity for each voxel, condition and subject was modelled using a canonical haemodynamic response function plus temporal and dispersion derivatives. For each individual, the effect of switch and repeat conditions was determined (switch>repeat, and repeat>switch). Statistical parametric maps of the t-statistic (SPM{t}) were generated for each subject and the contrast images were stored. In a second level random-effects analysis, these images were then combined in a one-sample t-test model was used (a p<0.001 at the voxel level was applied in all cases). The surviving activated voxels were superimposed on high-resolution structural magnetic resonance (MR) scans of a standard brain [Montreal Neurological Institute (MNI)]. Anatomical identification was performed with reference to the Talairach Daemon Software (http://www.talairach.org/) and also via XjView8 (http://www.alivelearn.net/xjview/). In addition, the neuroanatomy atlases by Haines (2011), and Nolte and Angevine (2007) were consulted.

### TMS Methods

Before their participation a high-resolution structural MRI was obtained from each participant to rule out any brain abnormalities, to locate the targets for TMS, and to guide the stimulation during de TMS sessions. The MRI was done on a 1.5T scanner (Siemens Magnetom Essenza). Sequences obtained were FSPGR-T1 3D (180 1-mm-thick sagittal slices, TR = 500 milliseconds, TE = 50 milliseconds, 256×256 matrix, FOV = 24 cm), SE T1, FSE DP-T2, FLAIR, and diffusion.

All participants underwent three separate sessions in which they received one of the three stimulation conditions: TMS over prefrontal areas, TMS over parietal areas or sham stimulation positioning the coil tilted 90° over vertex (control condition). After each of the stimulation conditions the participants performed the task. The order of the sessions was counterbalanced across participants. The stimulation was performed using a Magstim Super Rapid 2 stimulator (Magstim Company Ltd., Whitland, U.K.) with a 70 mm figure of eight-coil and the position of the coil location during stimulation was guided by a combination of Brainsight (Rogue Research, Montreal Canada) frameless stereotaxic system and Polaris (Northern Digital, Waterloo, ON, Canada) infrared tracking system.

At the beginning of the first session, and before stimulation, active motor threshold (aMT) was individually determined in order to set the appropriate output TMS intensity for each participant. To calculate the aMT, the coil was placed tangentially over the participant’s right M1, with the handle positioned 45° backwards. The coil was repositioned until the hot-spot of the left hand first dorsal interosseous (FDI) was located. The aMT was defined as the minimum stimulator output that produced movement in 5 of 10 trials in the FDI when the muscle was gently contracted (approximately 20% of the maximum voluntary contraction). The mean aMT value for the group was 53.4±6.2% of the maximum stimulator output and the intensity applied was the 80% of aMT. After determining aMT, and before stimulation, participants performed practice trials of the experimental task.

Continuous Theta Burst Stimulation (cTBS) was used in order to induce transient decreases of local cortical excitability lasting beyond the stimulation patterns (Huang et al., 2009; Huang et al., 2011). The cTBS protocol consists on a total of 600 TMS pulses grouped in trains of three pulses at 50 Hz, with each train being repeated every 200 milliseconds (5 Hz). The stimulation lasted 40 s in total and was delivered at 80% of the aMT of each participant (Huang et al., 2009; Rossi et al., 2009). After the cTBS, participants completed the task in a PC with a 17-inch monitor, the response buttons were letters M and Z of the keyboard. As part of the security protocol, in each session participants completed Mini Mental State Examination and Beck Depression Inventory before and after TMS to control for any changes in mood or cognitive functioning.

Targeted cortical areas were localized individually using the high-resolution MRI and the coordinates obtained from the previous fMRI study. MNI coordinates corresponding to maximum activation during the task performance were x=48, y=34, z=22 for prefrontal area, and x=42, Y=-54, z=58 for parietal area. Since these coordinates correspond to the anterior portion of the IFJ (aIFJ) and the IPS, coordinates were adjusted for individual anatomy to the closest gyrus area to the aIFJ and the IPS using Brainsight. For sham stimulation, the coil was positioned on vertex and tilted 90° to mimic the sound of the stimulation patterns used in the active conditions.

## Results

### fMRI Results

#### Accuracy

Subjects performed the task efficiently during the fMRI session and committed less than 17% errors. The average percentage of correct trials was 94.9%.

The Student *t* test on the percentage of correct responses during switch and repeat trials revealed the presence of marginally significant differences (t(19)=-1.8; p< 0.088) in line with a switch cost phenomenon where participants achieved less correct trials after switch cues as compared to repeat cues (94.5% versus 96%, respectively).

#### Reaction Times

The Student *t* test on RTs to switch and repeat trials revealed the presence of significant differences (t(19)=3.71; p< 0.001) indicating behavioural switch cost with an increase in RTs during switch as compared to repeat trials (737.2±34.9 versus 697.7±29.4, respectively).

#### fMRI activation results

Functional images were analysed by SPM12 using a general linear model applied at each voxel across the whole brain. We localized those brain areas that modulated their activity during the switch and repeat events. We focused on voxels for which the difference between the responses to the switch events and the repeat conditions was statistically significant. Results showed that cue-related brain activation in the switch > repeat contrast involved the right frontal, both left and right parietal lobes and in the cerebellum (see Table 1; Figure 2a). Target-related brain activation in the switch > repeat contrast involved a widespread number of brain areas within prefrontal, parietal, temporal, occipital lobes, and cerebellum (see Table 2, and Figure 2b).

**Table 1:** Locations and t-values of the main clusters of fMRI activity for the switch>repeat contrast during cues.

| Lobe | Region | BA | H | cluster | x | y | z | t-value |
| --- | --- | --- | --- | --- | --- | --- | --- | --- |
| Frontal | Inferior Frontal Junction anterior | 45 | R | 539 | 48 | 34 | 22 | 5.59 |
|  | Inferior Frontal Junction posterior | 44 | R |  | 50 | 10 | 24 | 5.37 |
|  | Superior Frontal Gyrus | 6 | R | 27 | 14 | 12 | 54 | 4.36 |
| Parietal | Intraparietal Sulcus | 7 | L | 137 | -10 | -76 | 60 | 5.04 |
|  | Intraparietal Sulcus | 40 | L | 15 | -32 | -80 | 48 | 4.38 |
|  | Intraparietal Sulcus | 7 | R | 553 | 8 | -74 | 48 | 4.97 |
|  | Intraparietal Sulcus | 40 | R |  | 42 | -54 | 58 | 4.95 |
| Cerebellum | Cerebellar Tonsil posterior |  | L | 161 | -22 | -64 | -32 | 7.57 |
|  | Cerebellar Tonsil posterior |  | R | 205 | 36 | -56 | -38 | 5.56 |

**Table 2:**
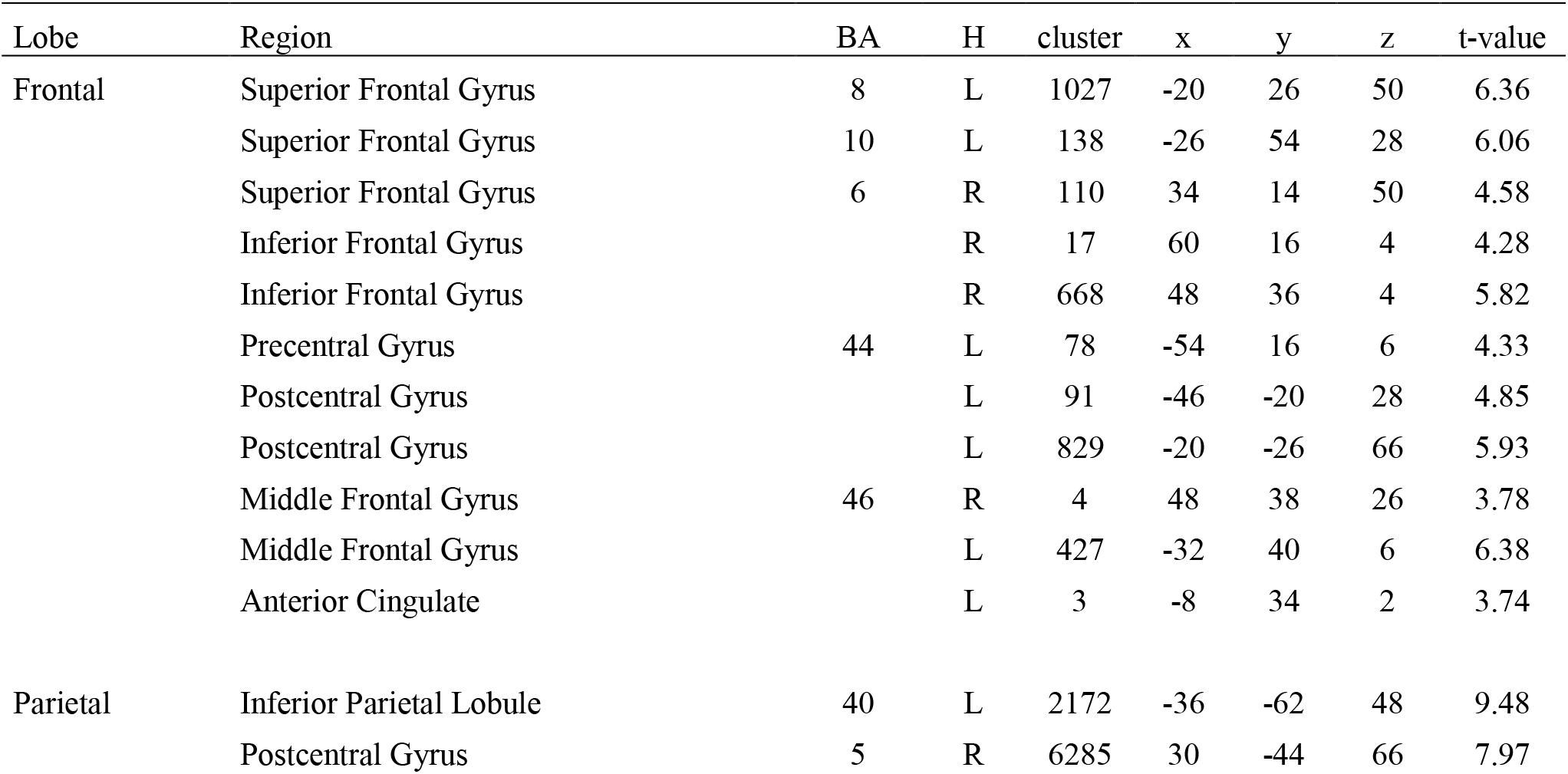
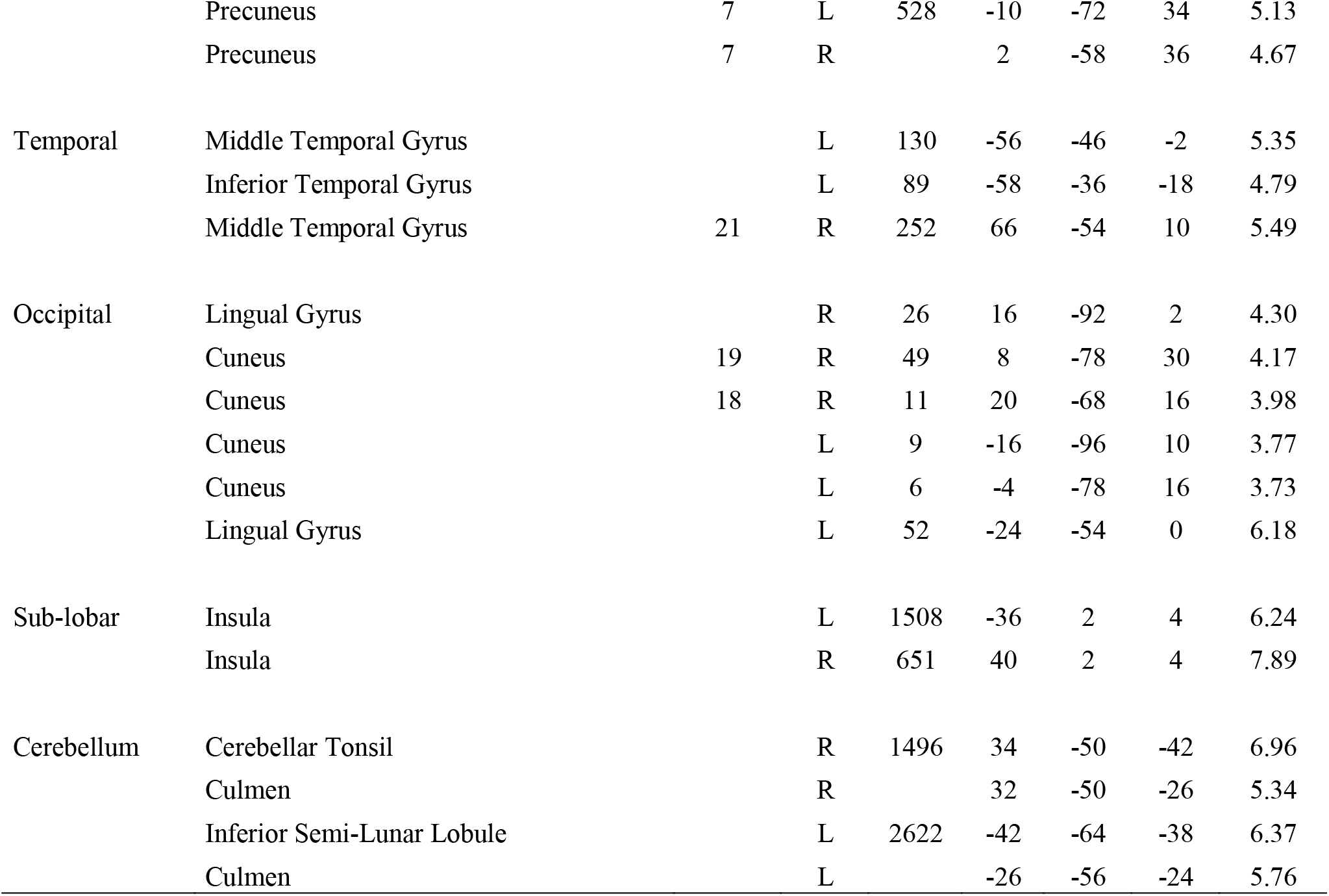
Locations and t-values of the main clusters of fMRI activity for the switch>repeat contrast during targets.

**Figure 2:**
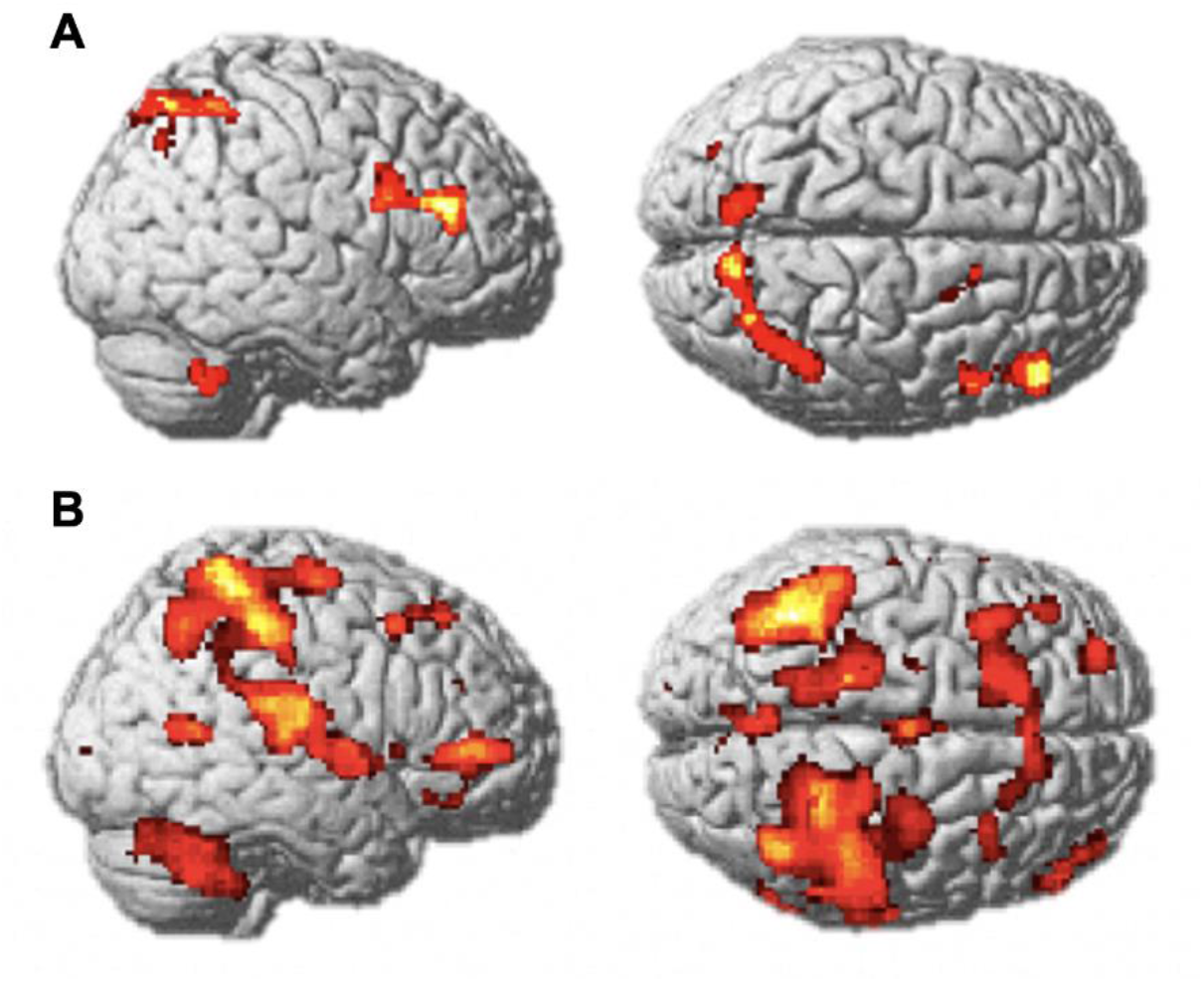
(a) fMRI group activation map showing activated brain regions when comparing cues in the switch>repeat contrast. Two main clusters of activations were identified at the anterior portion of the inferior frontal junction, and the intraparietal sulcus. (b) fMRI group activation map showing activated brain areas when comparing targets in the switch>repeat contrast. Activation was identified at a widespread network of brain areas within prefrontal, parietal, temporal, occipital lobes, and cerebellum. Images thresholded at p < 0.001. Neurological convention is followed (left side of the brain is shown on the left side of the figure).

### TMS results

#### Accuracy

Subjects performed the task efficiently during the TMS session and typically committed less than 13% errors across each of the three experimental conditions. The average percentage of correct trials across conditions was 95.8%.

The 3 × 2 repeated measures ANOVA on the percentage of correct responses using Area (aIFJ, IPS, and Sham stimulation) and Task (switch and repeat) as the factors revealed the presence of a main Task effect (F(1, 19)=10.1; p< 0.005), indicating a switch cost phenomenon with less correct trials after switch cues as compared to repeat cues (95,6% versus 97,7%, respectively). Neither the effect of Area (F(2, 38)=1.18; p= 0.889), nor the Area × Task interaction (F(2, 38)= 7.19; p= 0.494) reached significance.

#### Reaction Times

A 3 × 2 repeated measures ANOVA with RTs as the dependent variable was performed to compare the effect of TMS stimulation in the three different brain areas (aIFJ, IPS, and Sham) as a function of Task (switch and repeat). A significant main effect of Area was found (F(2, 38)= 5.94; p< 0.006) indicating that, overall, the stimulation of the aIFJ resulted in a significant increase of RTs across Task conditions as compared to Sham stimulation (p< 0.020), being the effect of IPS stimulation only marginal (p= 0.078), and with no differences in RTs between aIFJ and IPS conditions (p = 0.514). The presence of a main effect of Task (F(1, 19)=11.75; p< 0.003) confirmed results from accuracy analyses, regarding the presence of a significant switch cost effect when switch and repeat trials were compared (693.7±32.4 versus 657.8±26.7 milliseconds, respectively). More importantly was the presence of an Area × Task interaction (F(2, 38)= 4.25; p< 0.022). Post-hoc analyses of this interaction revealed that while the effect of the aIFJ stimulation disrupted RTs in both switch (p< 0.02) and repeat (p< 0.043) conditions as compared to Sham stimulation, the effect of the IPS stimulation had a specific negative impact over switch trials (p< 0.012) but not in repeat trials (p< 0.783). No differences were found when RTs to switch trials were compared between the aIFJ and the IPS stimulation (p= 0.921), or when they were compared in repeat trials (p= 0.267; see Figure 3).

**Figure 3:**
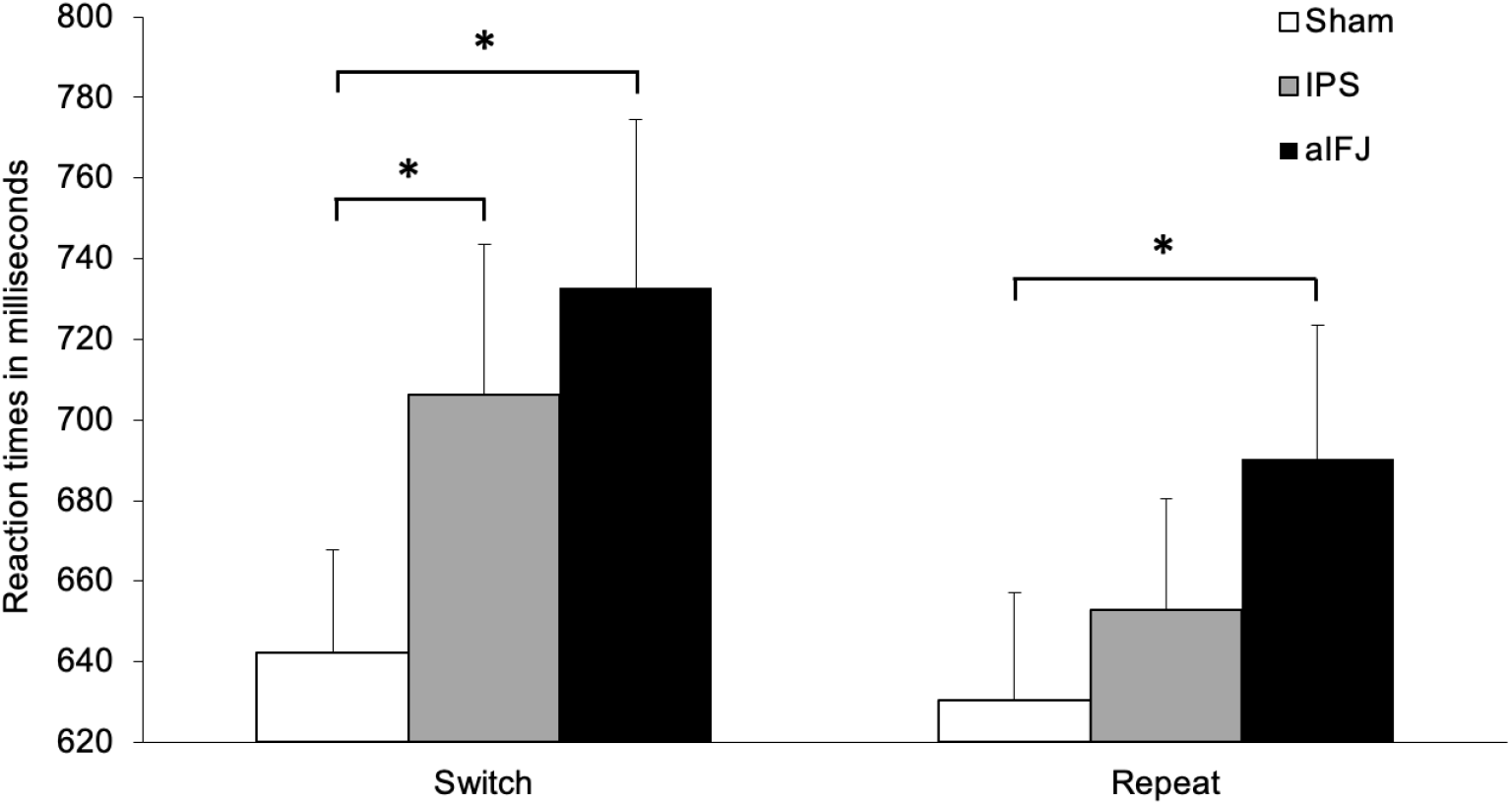
Reaction times during task switching performance in the TMS experiment. Mean reaction times in milliseconds (± S.E.M) for the task switch and the task repeat conditions, across the three conditions of TMS: aIFJ (stimulation of the anterior portion of the Inferior Frontal Junction), IPS (stimulation of the intraparietal sulcus), or Sham stimulation.

## Discussion

The aim of the present fMRI-TMS combined study was investigating the disruptive effect of TMS over specific prefrontal and parietal cortices within a FPN of brain regions previously related to preparatory task-switching. As expected, the results of the fMRI study revealed a behavioural switch-cost with an increase in RTs during switch as compared to repeat trials. This behavioural cost was accompanied by an increased activation in the IFJ (anterior and posterior portions), and the IPS during switch compared to repeat cues (see Tables 1, and Figure 2a). These findings are coherent with a large body of evidence suggesting the involvement of a FPN in preparatory process during task-switching paradigms (Dove et al., 2000; Gruber et al., 2006; Jamadar et al., 2010; Ruge et al., 2013). Interestingly, the same switch versus repeat contrast did not reveal activation differences in these brain regions during target processing. In this regard, switch-specific activation during target processing included a widespread network of brain areas within prefrontal, parietal, temporal, occipital lobes, and cerebellum (see Table 2, and Figure 2b). These results led us to select the aIFJ, and the IPS as the brain regions for the TMS study, since they reached the higher activation in prefrontal and parietal areas, respectively (see Table 1).

TMS results revealed that reducing excitability in the aIFJ, and IPS had dissociable behavioural effects. TMS over the aIFJ disrupted performance in both switch and repeat trails in terms of delayed responses as compared to Sham condition. In contrast, TMS over the IPS selectively interfered performance in switch trials (see Figure 3). These results suggest that the IPS would be related to the implementation of a switch-specific process while the aIFJ would be involved in a more general switch-unspecific process of task preparation. Functional neuroimaging and neurophysiological evidences agree with the present results showing that preparatory task-switching would involve both general and specific processes associated to certain prefrontal and parietal areas, respectively. For instance, using fMRI during cued task-switching blocks against single-task blocks, Braver et al. (2003) found that, while the activity in inferior prefrontal areas was selectively associated with mixing costs (i.e., the extra time it takes to perform two tasks simultaneously), only IPS activity was related to the magnitude of the switch cost. In agreement with the present results, the authors interpreted that prefrontal areas would support sustained components of cognitive control, such as the maintenance demands associated with keeping multiple task-sets at a relatively high level of activation, common to both switch and repeat trials. In contrast, parietal areas would support a more transient control process associated with task-switching, such as the updating of goals or the linking of task cues to their appropriate stimulus-response mapping. In another study integrating event related potentials and fMRI data, Jamadar et al. (2010) found that informative task-switch cues (switch or repeat) evoked larger amplitudes in an early cue-locked positive component as compared to cues providing no information about the next task (non-informative cues). Importantly, this amplitude modulation correlated with fMRI activity in lateral prefrontal cortex (cued informative versus cue non-informative contrast), consistent with a general goal activation process. The authors also reported a cue-locked late positivity that was larger for informatively cued switch versus repeat cues, whose amplitude correlated with fMRI activity in the posterior parietal cortex (cued switch versus cue repeat contrast), compatible with a stimulus-response rule activation process. These findings would indicate that lateral prefrontal brain regions seem to relate to general preparation processes, instead of specific switching operations. Also, in another ERP study, Barceló et al. (2008) found enhanced fronto-centrally distributed cue-locked P3 activity in response to informative cues in a task-switching paradigm irrespective of switch or repeat demands, compared to analogous irrelevant cues in an oddball control task. However, only a late cue-locked P3 component in parietal areas was specifically modulated in response to switch versus repeat equiprobable cues. The authors suggested an association between the fronto-centrally P3 and a general control mechanism related to preparatory process in frontal areas, and between the posterior P3 and a switch-specific reconfiguration process for stimulus-response mapping in parietal areas. Taken together, the above-mentioned evidences and the present results support a multi-component view of anticipatory preparation in task-switching (Ruge et al., 2013). It could be that a general preparation process, such as the retrieval of cue information for goal identification, necessarily during both switch and repeat trials, will recruit the aIFJ. In this vein, areas in the vicinity of the aIFJ reported here have been widely related to different components of memory retrieval. For instance, Nyhus and Badre (2015), associate BA 45 (also referred as mid-ventrolateral prefrontal cortex) to post-retrieval selection needed both to solve competition among multiple memory representations and to align remembered information with current task goals. Also, the application of inhibitory TMS over mid-ventrolateral prefrontal cortex selectively diminishes recall for episodic details in an old/new memory-discrimination task, and lead the authors to attribute a causal role of this area in goal-directed retrieval (Wais et al., 2018). On the other hand, a switch-specific preparation processes such as the translation of abstract task goals into specific action rules or stimulus-response maps for task implementation necessarily during switch trials only, seemed to recruit the IPS. The IPS has been largely related to response selection processes, or more precisely, to stimulus-response mapping. Worringer et al. (2019) interpreted that stronger IPS activation in switch versus repeat trials may reflect the controlled mapping of a given stimulus to the response that is adequate according to the current task-set. IPS projections to ventral premotor areas support the role of this areas in action planning and reorienting of motor attention (Binkofski et al., 1998; Rushworth et al., 2003). The specific area of parietal activation during task-switching may reflect variations based on specific task stimulus attributes, with numerical tasks like the one used here (i.e., >5/<5, and odd/even) activating IPS within the right hemisphere (Shi et al., 2014).

It has to be noted that the present findings differ from a previous TMS study using a task-switching paradigm (Muhle-Karbe et al., 2014). As mentioned above, these authors found that the application of inhibitory TMS over both the IFJ and IPS interfered performance over switch trials only, suggesting that both prefrontal and parietal regions would implement switch-specific preparation processes. On the contrary, the present results revealed that while inhibitory TMS over the aIFJ interference performance in both switch and repeat trials, TMS over IPS produced interference only in switch trials. Although apparently contradictory, a fine-grained anatomical analysis revealed that two different regions within the IFJ were stimulated in these two studies. Particularly, the prefrontal brain area stimulated by Muhle-Karbe et al. (2014) (MNI: -40, 4, 32, BA 44) was more posterior to the aIFJ (MNI: 48, 34, 22, BA 45) reported in the present study (note that both areas were activated in the present fMRI study). Such a distinction between anterior and posterior portions within the IFJ is coherent with current models suggesting the existence of a hierarchical rostro-caudal organization in lateral prefrontal cortex based in the abstractness of action representations (Koechlin and Summerfield, 2007; Badre and D’Esposito, 2009; O’Relly, 2010). On the one hand, the aIFJ would be involved in a general preparatory process such as the retrieval of episodic information for goal selection (Buckner, 2003). This operation would take place at a relatively high level of abstraction any time a cue is presented, and irrespective of switching or repeating task. On the other hand, the pIFJ would be involved in a more specific preparatory process such as the updating of task-goals representations (Derrfuss et al., 2005; Brass et al., 2005). Following Muhle-Karbe et al. (2014) while the pIFJ (BA 44) would provide an initial representation of task goals that needs to be selected during the next switch trial at a more abstract level, the IPS might specify the resulting stimulus-response mappings, thereby providing a more action-related task representation (Brass and von Cramon, 2004; Brass et al., 2005). These interpretations fit well with the functional fractionation between episodic and contextual control as proposed by the hierarchical model about the architecture of cognitive control in prefrontal cortex (Koechlin & Summerfield, 2007).

In conclusion, the pattern of TMS results observed supports a model of task-switching preparation that involves both general and specific reconfiguration processes. Particularly, the present results reveal for the first time that inhibitory stimulation in the aIFJ (BA 45) may have caused interference in the retrieval of episodic information for goal selection during task-switching. This interpretation is coherent with current data about the role of aIFJ in memory retrieval under high control demands (Buckner, 2003). In addition, our results confirm the purported role of the IPS in more specific mechanisms of goal setting and implementation (Binkofski et al., 1998; Rushworth et al., 2003). Further research needs to clarify the pinpointed dissociation between the anterior and posterior regions of IFJ. Combining fMRI and TMS data might be critical to provide a more complete picture about the complex interaction between brain areas within the frontoparietal network and cognitive control operations during task-switching.

## Acknowledgements

Supported by the Spanish Ministry of Science and Innovation, Instituto de Salud Carlos III (ISCIII), CIBER -Consorcio Centro de Investigación Biomédica en Red- (CB/07/09/0023), financed by the European Union, ERDF Funds from the European Commission, “A way of making Europe”, (PI1001061), Madrid Regional Government (B2017/BMD-3740 AGES-CM-2), European Union Structural Funds, European Union Seventh Framework Program, European Union H2020 Program under the Innovative Medicines Initiative 2 Joint Undertaking: Project PRISM-2 (Grant agreement No.101034377), Project AIMS-2-TRIALS (Grant agreement No 777394), Horizon Europe, the National Institute of Mental Health of the National Institutes of Health under Award Number 1U01MH124639-01 (Project ProNET) and Award Number 5P50MH115846-03 (project FEP-CAUSAL), Fundación Familia Alonso, and Fundación Alicia Koplowitz.

